# SpineS: An interactive time-series analysis software for dendritic spines

**DOI:** 10.1101/2020.09.12.294546

**Authors:** Ali Özgür Argunşah, Ertunç Erdil, Muhammad Usman Ghani, Yazmín Ramiro Cortés, Anna Felicity Hobbiss, Theofanis Karayannis, Müjdat Çetin, Inbal Israely, Devrim Ünay

## Abstract

Live fluorescence imaging has shown the dynamic nature of dendritic spines, with changes in shape occurring both during development and in response to activity. The structure of a dendritic spine positively correlates with its functional efficacy. Learning and memory studies have shown that great deal of the information stored by a neuron is contained in the synapses. High precision tracking of synaptic structures can give hints about the dynamic nature of memory and help us to understand how memories evolve both in biological and artificial neural networks. Experiments that aim to investigate the dynamics behind the structural changes of dendritic spines require the collection and analysis of large time-series datasets. In this paper, we present an open-source software called SpineS for the automatic longitudinal structural analysis of dendritic spines with additional features for manual intervention to ensure optimal analysis. Our extensive experimental analyses on multiple datasets demonstrate that SpineS can achieve a high-level performance on samples collected both by two-photon and confocal imaging systems.

## Introduction

The efficacy of excitatory synapses changes with development^1^, activity^2^ and learning^3^, and correlates with structural changes of dendritic spines^4–9^. Changes in efficacy and structure reflect activity at a synapse, and can impact subsequent information transmission between inputs across the dendritic arbor^10–12^. Understanding how such changes are physically maintained in the neuron is key to elucidating the mechanisms by which information is stored in the brain. Activity-dependent structural changes at spines can last from minutes to days, and are experimentally visualized through multi time point sampling of z-stack images, often collected over many hours. For example, in an experiment that addresses structural plasticity mechanisms along a dendritic branch using fluorescence imaging, depending on the image acquisition conditions and the type of the neuron that is being imaged, hundreds of spines can be assessed. In a longitudinal study, the total number of spines to be analyzed can reach up to thousands and analyzing such a dataset manually is tedious and time consuming.

The focus of investigation into structural changes of dendritic spine features has thus far been centered on changes within the spine head volume, spine neck length and density of spines at a dendritic segment. Two methods have emerged in the field for the estimation of spine volume, namely integrated fluorescence intensity (IFI) and full-width at half maximum (FWHM^)8,13^. The IFI method sums all the fluorescence values within a region of interest (ROI) drawn around the spine of interest^7, 14, 15^. In the FWHM method, an intensity profile over a line perpendicular to the spine neck and passing through the spine head center is used to fit a Gaussian. The half maximum value of the estimated Gaussian fit is used to estimate the diameter of a hypothetical sphere representing the spine head. IFI is sensitive to fluctuations of fluorescence intensity caused by the imaging system, which can be dramatic; whereas FWHM suffers from over or under estimation if the spine head is not a perfect sphere which often is the case^16^. In order to overcome any intensity variation due to changes in expression of fluorescent proteins that might occur during imaging, the IFI volume estimation can be corrected by normalizing with the fluorescence intensity at the nearby dendrite after background subtraction^15, 17^.

Further studies have revealed that spine neck features such as width and length are also important structural modifications that correlate with activity^9, 18^. It has been shown that spine neck gets shorter and thicker as spine head gets bigger following long-term potentiation (LTP).

In this paper, we present an open-source Matlab-based software called SpineS that allows: (1) automatic registration of dendritic arbor images collected at consecutive time points to correct for possible spatial displacements that could happen during image acquisition, (2) automatic segmentation of spine heads, (3) calculation of spine head volumes using IFI and spine neck lengths from two-photon laser scanning microscopy (2PLSM) and confocal laser scanning microscopy (CLSM) image stacks. SpineS also provides tools for reviewing and correcting dendrite and spine head segmentations and spine neck paths, as well as a tool to manually estimate FWHM-based spine head volumes. We choose Matlab since it is widely used by biologist in image analysis task. The Matlab toolboxes that are necessary to run SpineS are Curve fitting, Image processing, Signal processing, and Statistics. In this paper we explain both the front and back-end inner workings of SpineS as well as providing extensive performance analysis on multiple datasets collected using different imaging systems to address different biological questions from multiple laboratories.

There have been various open-source software packages developed for dendritic spine analysis during the last decades, most of which are no longer available. Neuron-IQ^19^ was another Matlab-based software that is no longer downloadable. NeuronStudio^20^ was incorporated into the commercial software Neurolucida^21^. SpineJ^22^ is a recently released good alternative for super-resolution microscopy image stacks but whether it would work for CLSM or 2PLSM images is an open question and it does not offer time-series analysis. 2dSpAn^23^ and its 3D version 3dSpAn^24, 25^ are two other alternative available software packages for morphological analysis of dendritic spines. Although these software packages are providing good results, they also do not offer longitudinal analysis and do not allow for manual correction, and either report spine head and neck volume together as one entity or the total length of the spine head from the tip to base, neither of which have reported biological correlates. However an advantage is that they do not require Matlab to run as our software does.

## SpineS graphical user interface

The SpineS GUI includes all the necessary components to analyze spines on a dendritic branch (see Figure 1). In this section, we introduce these components and give the necessary instructions to conduct analysis with SpineS (also see Supplementary Figure S1).

**Figure 1.**
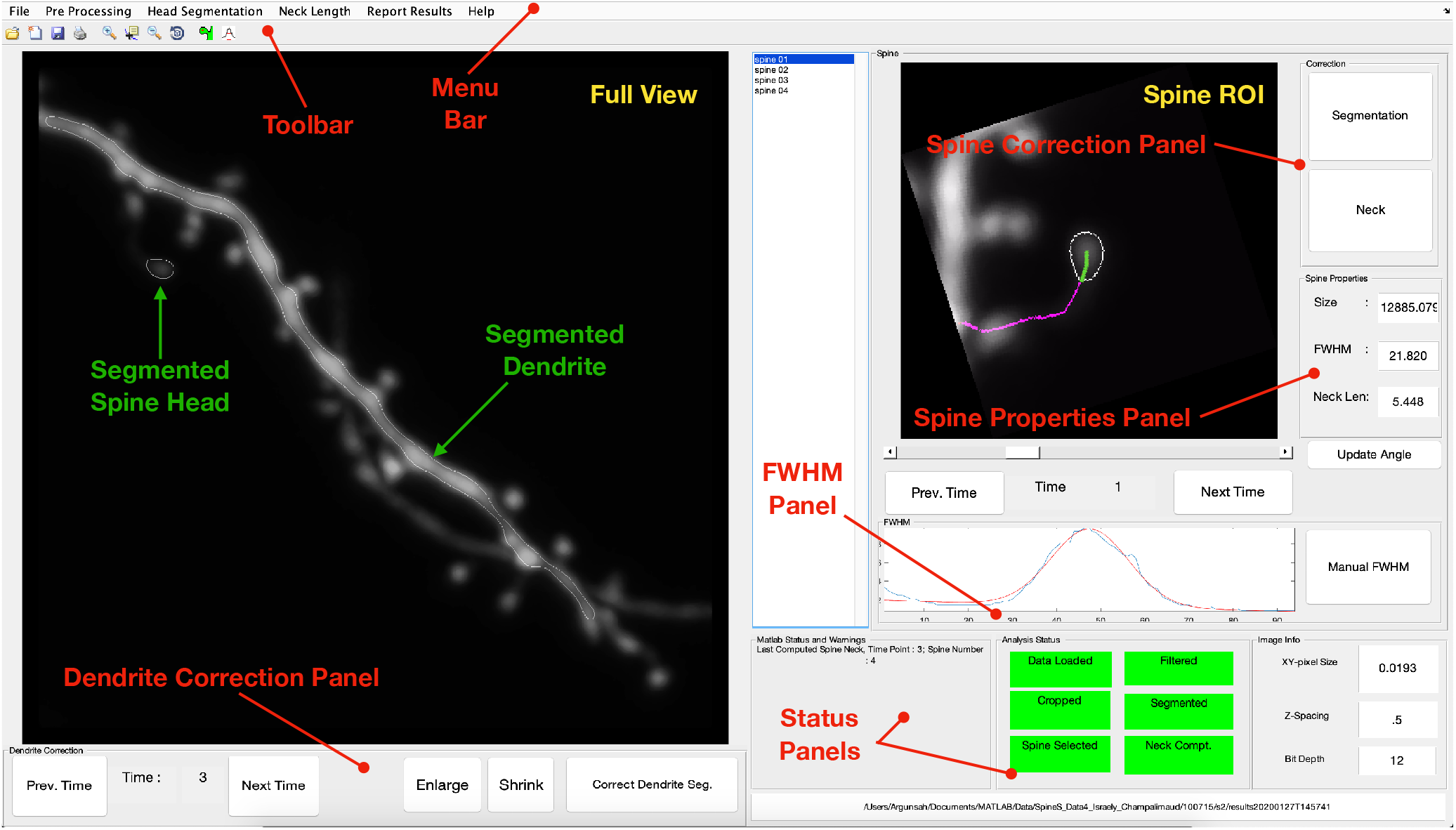
SpineS GUI allows automatic analysis of dendritic spines in combination with interactive manual correction tools. The *Full View* pane shows a maximum intensity projection (MIP) of the first time point of the dendrite to be analyzed after loading the data. Spines to be analyzed are selected by the user only at the first time point by clicking on the spine centers. SpineS automatically analyzes the selected spines for all time points.

### Steps to analyse a dataset

The first step that is carried out by the SpineS software is loading the data (Figure 2A). Since different laboratories use different imaging systems and image formats, data specifications can be very different. In SpineS, we use the bio-formats library provided by the open microscopy environment project to support all microscopy platforms^26, 27^. The software computes the maximum intensity projection (MIP) of the first time point to display the image for spine selection by the user as shown in Figure 1. SpineS analyzes only one dendritic branch at a time, hence if there are multiple dendrites in the view, the dendrite of interest should be cropped as the first preprocessing step (Figure 2B).

**Figure 2.**
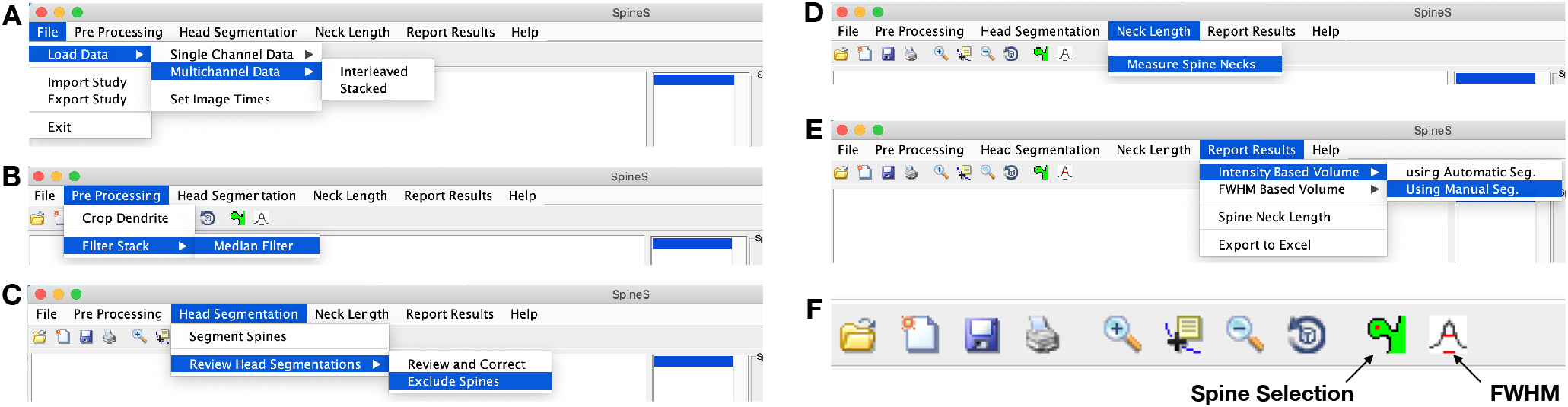
SpineS GUI Menu Bar (A) Single or multichannel times series data can be loaded either using the open microscopy bioformats plugin or directly from folders containing .tif image stacks. (B) If there are multiple dendrites, the dendrite of interest should be cropped manually. Images are filtered using a median filter with a user-defined filter size and images in all time points are registered afterwards. (C) The head segmentation tab includes the *Segment Spines* option to start the segmentation process and global correction as well as *Exclude Spines* from the analysis in case of erroneous spine segmentation. (D) After the dendrite and spine head segmentations are completed and corrections are performed, *Measure Spine Neck* function can be executed to compute spine neck paths. (E) When all previous steps are completed, the user can plot the results and export to an Excel format. (F) The toolbar includes spine selection and FWHM buttons in addition to classic Matlab tools.

**Figure 3.**
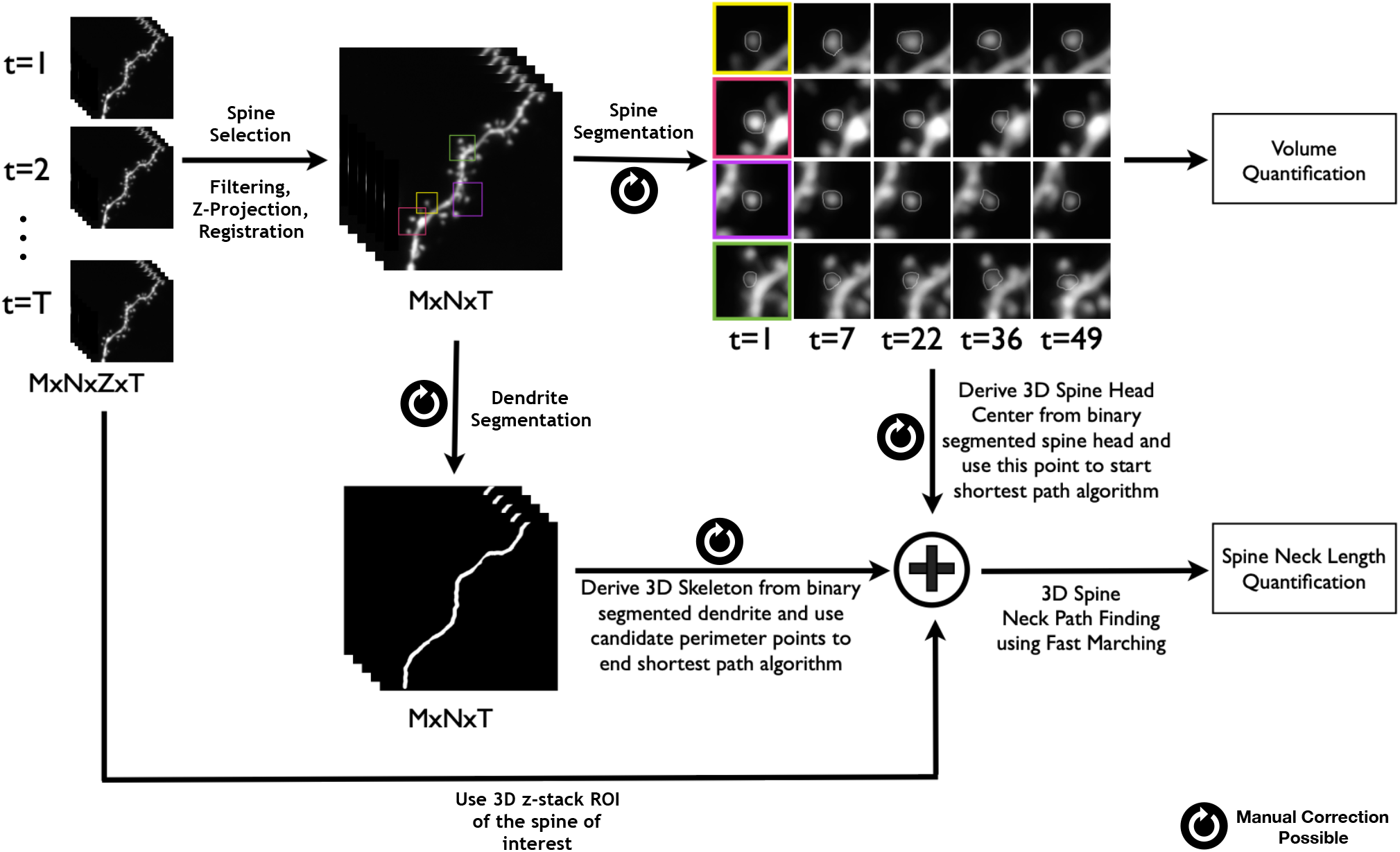
Workflow of SpineS. Z-stacks (MxNxZ) from multiple time points (T) are analyzed. Each z-stack is imported and registered. MIPs are then computed and filtered using median filtering. Dendritic spine heads are segmented using a watershed segmentation algorithm followed by a graph-based clustering. Each spine volume is estimated using integrated florescence intensity of spine intensities within the segmentation region. The intensities in the region are normalized using the median florescence intensity of the dendrite at the corresponding time point after background fluorescence subtraction. Neck paths from the spine head center to the closest geodesic point on the dendrite are computed using the fast-marching algorithm by imposing empirically set constraints.

Spines to be analyzed should be selected by the user by clicking the spine head centers for the first time point using the *Spine Selection* icon (Figure 2F). Once the selection is completed, the *Median Filter* option under the *Filter Stack* applies a median filter to all MIP images in the time series and the filtered images are registered to the first image (Figure 2B). Following the automatic dendrite segmentation, *Dendrite Correction Panel* in Figure 1 allows the user to fine-tune the segmentation. Whilst editing the segmentation, it is possible to enlarge or shrink the entire segmented dendrite. The user can shrink or expand the boundaries via an active contour based segmentation^28^. For a more localised problem, the *Correct Dendrite Seg*. button allows the user to correct specific parts of the segmentation mask (see Supplementary Figure S2).

After ensuring a proper dendrite segmentation at every time point, spine head segmentation can be run (Figure 2C). The *Head Segmentation* tab also includes a *Review and Correct* button which should be run after the completion of the head segmentation process for a gross correction of all head segmentations. Once the automatic spine head segmentation is completed, the user can navigate through the time points to inspect the quality of the segmentations and refine them if necessary by using the *Segmentation* button in the *Correction* panel in Figure 1. Once pressed, this button will present a new image window with the segmentation in a draggable correction mesh (see Supplementary Figure S3).

After all head segmentation corrections are completed, the automatic spine neck path algorithm can be run (Figure 2D). Erroneous neck paths can be corrected using the *Neck* button in the *Correction* panel shown in Figure 1 by providing a new base point on the dendrite.

If a FWHM-based volume estimation is needed, the spine orientation should be changed by using the slider below the *Spine ROI* panel such that the spine head is oriented vertically as shown in Figure 1-Spine ROI. When an orientation is modified at an earlier time point, all consecutive time points for that spine are registered automatically. FWHM requires the selection of a rectangular region including spine head center and surrounding background. This process can be started with the FWHM icon (Figure 2F).

Spine properties such as IFI-based head volume, FWHM-based head volume and neck length are shown in the spine properties panel (see Figure 1) after the completion of the necessary steps. Throughout the analysis, progress can be followed in the *Analysis Status* panel as the background color of each process turns from red to green upon completion (see Figure 1). The analysis results can be exported to MS Excel and figures will be created and displayed upon the completion of the analysis (Figure 2E).

## Methods

In this section, we present the image processing methods that were used to develop SpineS.

### Registration

During the imaging of dendritic segments, mechanical movements of the sample or the dynamic nature of dendrites and spines can cause movements in the images. In order to correct for such translational movements, we apply a two step registration procedure.

#### Global Registration

Global registration aligns two consecutive image stacks at sub-pixel precision using a discrete Fourier transform (DFT) based algorithm^29^. This algorithm uses selective upsampling by a matrix-multiply DFT to reduce computation time and memory while preserving accuracy.

#### Local Registration

Although global registration broadly aligns two consecutive time points well, some local misalignments may still remain due to orientation and shape changes of dendritic spines. As a result, the spine of interest might slightly shift from the center of the ROI. In SpineS, we force the spine of interest to be at the center of the ROI using local registration to avoid analyzing an incorrect spine that might appear in the ROI after the global registration. We achieve this using a Normalized Mutual Information (NMI)^30^ based local registration method. In this method, after global registration we shift the ROI within a small neighborhood, and measure the NMI with the ROI in the previous time point. Finally, we take the ROI with the highest NMI and use it in the analysis.

### Dendrite Segmentation

Dendrite segmentation starts by filtering the MIP image using a 2D-median filter and binarizing via Otsu thresholding^31^ to get a rough segmentation of the dendritic branch including spines. The medial axis of the dendrite is computed by applying a fast marching distance transform^32^ on the dendritic segment between the two ends of the dendrite. A locally adaptive sized disk-shaped structuring element around the medial axis of the dendrite is applied to remove the connected spines. Additional refinement is performed by assuming that the dendrite diameter remains constant within the local field of view.

### Spine Head Segmentation

We use a multilevel algorithm for spine head segmentation^33^. The user clicks in the centre of every spine at the first time point to start the analysis. In SpineS, the head segmentation consists of multiple steps to ensure optimal results. The initial segmentation in the pipeline is performed using a watershed algorithm^34^. Watershed segmentation requires a seed point to start and a boundary to determine where to stop the segmentation. When determining the seed points, we use the assumption that the spine head contains a maxima region which we extract using a morphological transformation^35^. We feed the maxima regions as seeds to the watershed to start the segmentation. We determine the boundaries of the segmentation using Otsu’s thresholding^31^. Watershed segmentation usually finds larger boundaries than manual segmentation by experts. We therefore incorporate a second level of segmentation algorithm. This takes each previously found component in the region of interest and refines it using a modified version of a graph theoretic algorithm for arbitrary shape detection^36^. This algorithm performs segmentation inside the region found by watershed which generates multiple clusters. These clusters are then combined using hierarchical clustering to obtain *k* = 10 number of clusters. These clusters are formed as quasi-concentric connected components. We exclude the outermost clusters and use those remaining as the final segmentation.

### Finding the Spine Neck Path

Neck length computation is a challenging task due to spine shape variations and neck motility. We begin computing the neck length by eroding each slice of the dendritic branch image with a disk-shaped structuring element to reduce spurious paths. Then, a multi-stencil fast marching (MSFM) method^32, 37^ is applied to compute the 3D distance map using the spine head centre as the source point. The Runge-Kutta algorithm^38^ is applied on the 3D distance map to compute the shortest paths (geodesic) from *N* points on dendrite perimeter to the spine head centre. These *N* points are selected by finding the *N* nearest points from the spine head centre to dendrite perimeter using the Euclidean distance as a metric.

The crucial final step is the selection of the correct neck path. A simple approach would be to select the path with the minimum length (Eq. 1), but that would be suboptimal due to the motile and sometimes non-linear nature of spine necks. Therefore, using the mere path length as a constraint is not enough. We introduced two additional constraints to select the path with best geodesic approximation: Path complexity (Eq. 2) (L1-norm of path derivatives), and path smoothness (Eq. 3) (L1-norm of image intensities along the path). We select the neck path that collectively has the lowest value for these three constraints (Eq. 4).

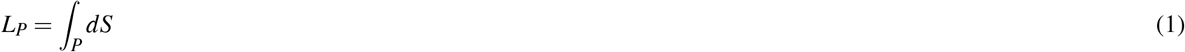

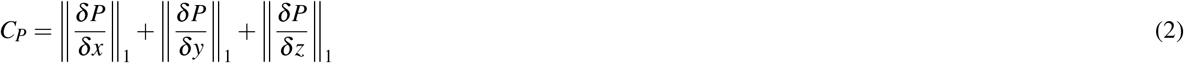

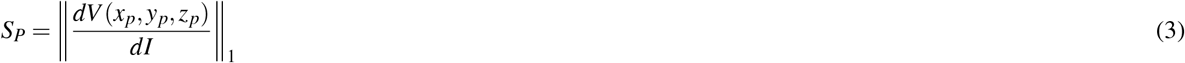

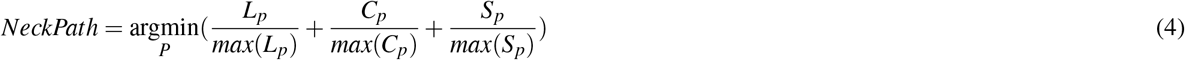

To compute the neck path, we remove the neck path passing inside the segmented spine head and dendrite borders.

An example spine neck path is seen in Figure 4. The green path in Figure 4B is the path of the computed spine neck that is inside the segmented head, hence not used to calculate the neck length. The magenta part shows the remaining spine neck path. Figure 4C shows that the spine base point is attached to the dendrite (green arrow head) but not to a nearby passing axon (blue arrow head).

**Figure 4.**
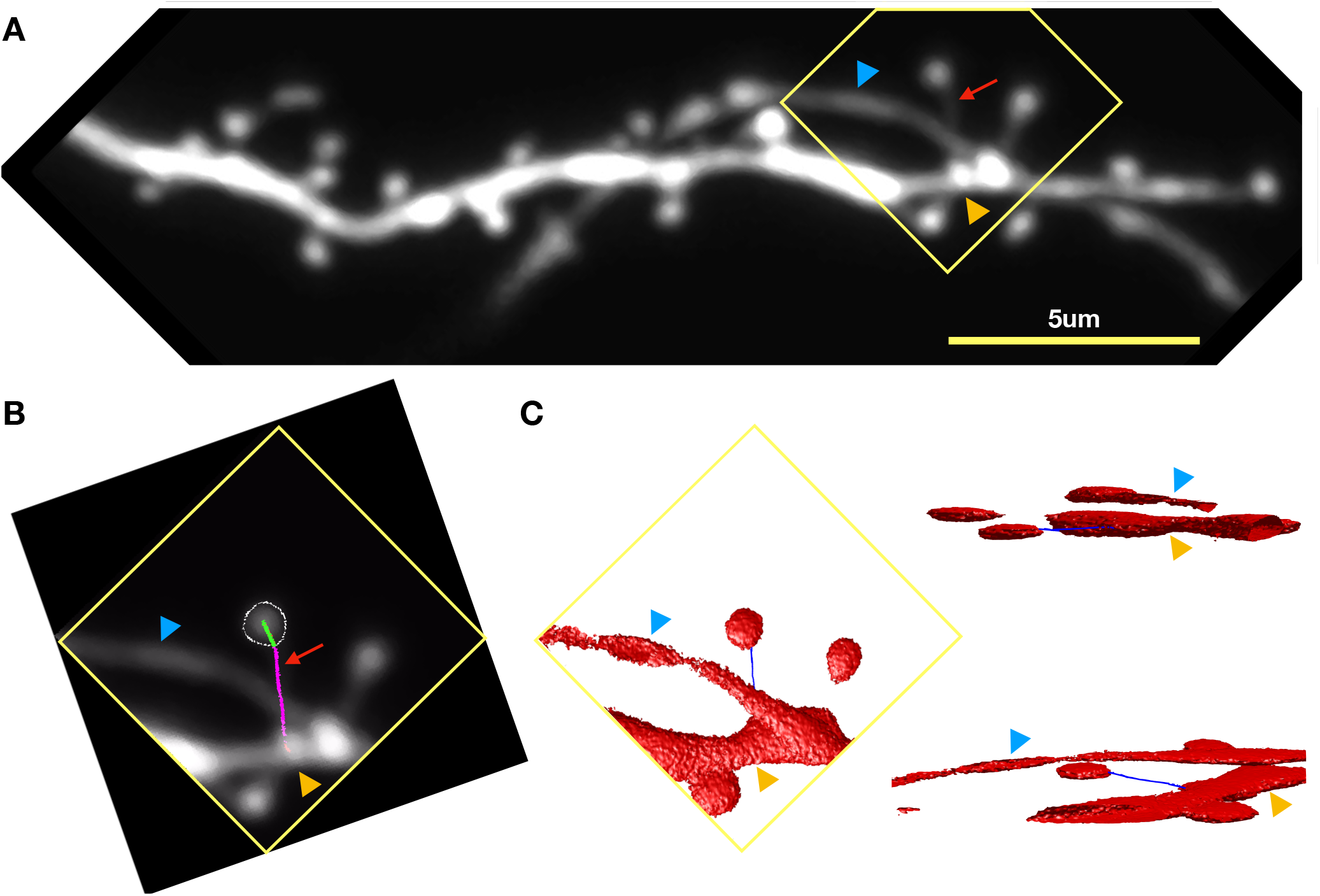
Finding spine neck path in 3D. (A) Two-photon microscopy image of a dendrite of a hippocampal CA1 pyramidal neuron in-vitro. The blue arrow indicates the nearby process. The orange arrow indicates the parent dendrite. Red arrow indicates the spine neck. Image contrast is increased to improve the visibility of the spine neck. (B) ROI in yellow square in A. The path from the parent dendrite to the spine head border is shown in magenta. The path goes through the spine head until spine head center is shown in green. The path is computed from the spine head center to dendrite center first. (C) 3-D rendered version of B showing the neck path is correctly traced from the spine center to the dendrite but not to the nearby process.

### Volume Estimation

#### Integrated fluorescence intensity

Intensity values within the segmented spine head are summed and the background fluorescence intensity (the minimum intensity value in the spine field of view (FOV) × area of segmented spine) is subtracted from the summed intensity value. Finally, an intensity normalization to the dendrite is necessary due to the relative changes in fluorescence level over time that might be caused by bleaching, movement or imaging laser power fluctuations^7, 15, 17^.

In earlier studies, summed intensities in the spine head were normalized using the highest intensity value of the dendrite portion closest to the spine. However, this approach might be problematic for two reasons. Firstly, if the collected images are saturated, the maximum recorded value is not a true representation of the brightness, thus the maximum intensity normalization produces misleading results. Secondly, since MIP images are collapsed along the z-axis, spines on this axis often appear as bright hotspots. If a nearby spine of interest is normalized to these hotspot-like points, one may end up normalizing to a floating reference. We believe this problem is either overlooked or solved but not reported in publications. In order to overcome this issue we use the median intensity value inside the segmented dendrite to normalize to, instead of the maximum intensity value. SpineS also reports the raw values for summed intensities, background fluorescence intensity and fluorescence intensity distribution within the segmented dendrite so that users can select the most relevant metric for their study.

#### FWHM-based volume estimation

It is often assumed that a spine has a spherical shape^5, 8, 39^. The FWHM method is used to find the radius, *r*, of a sphere that encloses the spine. Then, the spine volume is computed using the formula 4/3 *πr*^3^.

The fluorescence intensity profile is measured along a line parallel to the postsynaptic density (PSD), the active zone of the synapse normally located on the opposite side to the spine neck. A Gaussian curve is fitted to this intensity profile. Finally, the FWHM value of the Gaussian curve becomes the diameter (*R*) of the sphere. A benefit of using a FWHM method is that it is invariant to changes in fluorescence intensity. However there are some weaknesses of a FWHM-based volume estimation. This approach produces accurate results if the spine shape is close to spherical. However, most of the times, dendritic spines are non-spherical. Moreover, small variations in the rotation angle may change the estimated spine volume significantly. It is not easy to determine where the PSD is (to draw the fluorescence intensity profile) in MIP image, hence it is a challenge to find the best orientation automatically.

## Results

### Fluorescence Intensity invariant performance in spine segmentation

Since our IFI based volume estimation depends on the quality of the spine head segmentations, we wanted to see if significant fluorescence changes affect the segmentation performance. We collected a dataset by imaging the same dendritic segment using different laser intensities. To ensure that no significant biological remodeling occurred either at the dendrite or spines over the course of the in-vitro experiment we acutely added tetrodotoxin (TTX) into the artificial cerebrospinal fluid (ACSF) used to perfuse the neuron during the imaging session. It has been shown that application of TTX changes spine volumes only upon prolonged application^40^. We subsequently analyzed spines to investigate if the segmentation performance and resulting IFI volumes were sensitive to different levels of fluorescence. Figure 5A-D shows MIP images and segmented spines of the same dendritic segment imaged using four different imaging laser powers. The segmentation of spines was not affected by changes in fluorescence intensity. The average IFI drops as the imaging laser power decreases (see Figure 5E); however normalization to the dendrite recovers the expected volume invariance of the spines (see Figure 5F). As mentioned earlier, spine head width which is estimated using FWHM is not affected by intensity fluctuations and shows the linear trend without normalization (see Figure 5G).

**Figure 5.**
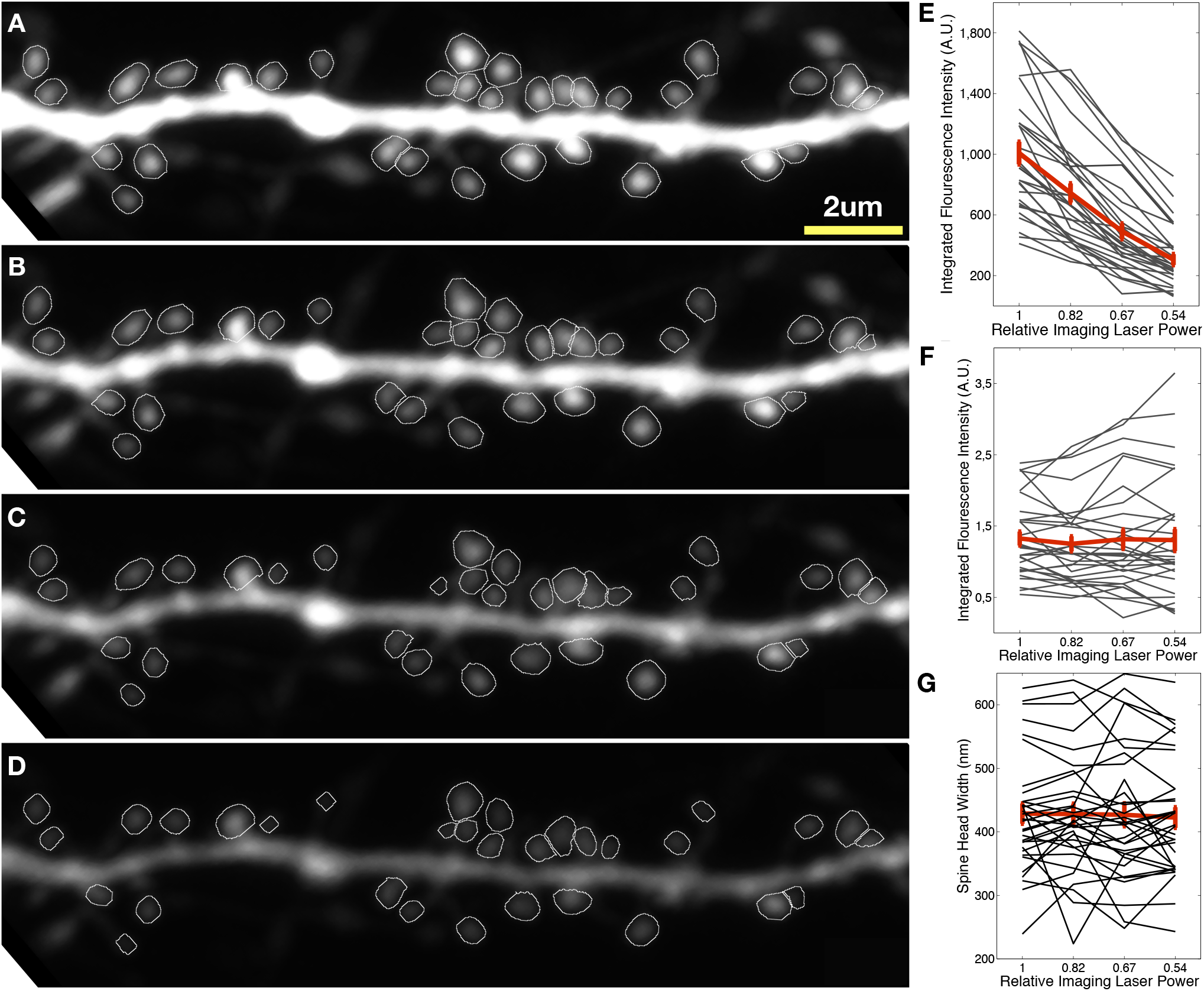
Maximum intensity projection of two-photon microscopy images collected using different imaging laser powers (A-D) as the tissue was perfused with ACSF containing TTX. (E) Individual (gray) and average (red) intensity based volume estimation of segmented spines. (F) Same as (E) after normalization with median intensity value at the dendrite (G) FWHM estimated spine head widths.

### Comparison with Ground Truth

We compared the normalized volumes of 27 spines from 9 different dendritic segments collected for 90 minutes at 5 minute intervals with data collected after analysis by human experts. It has been shown that application of the mGluR1-5 agonist (S)-3,5-Dihydroxyphenylglycine (DHPG) induces LTD at dendritic spines^41^. We analyzed this dataset using SpineS and compared the results of the software with those obtained by two experts in this domain, each manually performing one of the two spine volume estimation methods explained earlier - manual segmentation for IFI estimation and manual spine alignment for FMWH estimation with a tool designed for ImageJ. For comparison, we used the symmetric mean absolute percentage error (*sMAPE*) based similarity score (SS). The *sMAPE* is a robust measure for trend comparisons in time series data:

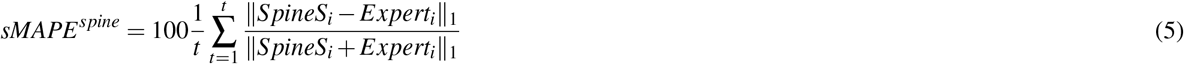

where, *t* represents time, *SpineS_i_* and *Expert_i_* represents the volume estimations of SpineS and Expert at time point *i*, respectively.

Using *sMAPE*, we compute the similarity score as follows:

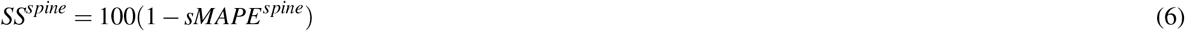

Comparisons of normalized volume results derived after SpineS segmentation, the manual segmentation and the manual FWHM estimation are given in Supplementary Table 1. Similarity scores of estimated intensity-based volumes from manually segmented spine-heads and SpineS output suggest that automatic segmentation yields very similar (*μ* = 90.28%; *σ* = 5.83%) spine head segmentations to the experts’, since both used an intensity-based volume estimation method. We also compared our results with the manual FWHM volume estimation results to see how comparable the two volume estimation methods used in the field are. In Figure 6 the caption indicates the similarity score of the two different methods for a given spine. In Figure 6F we see that on average, all 3 methods (the proposed tool plus the experts) converge to the same statistical distribution (all pairwise t-test *p* < 0.01). In Figure 6 we also present five examples of spine volume trends out of the 29 spines we used for the comparisons.

**Figure 6.**
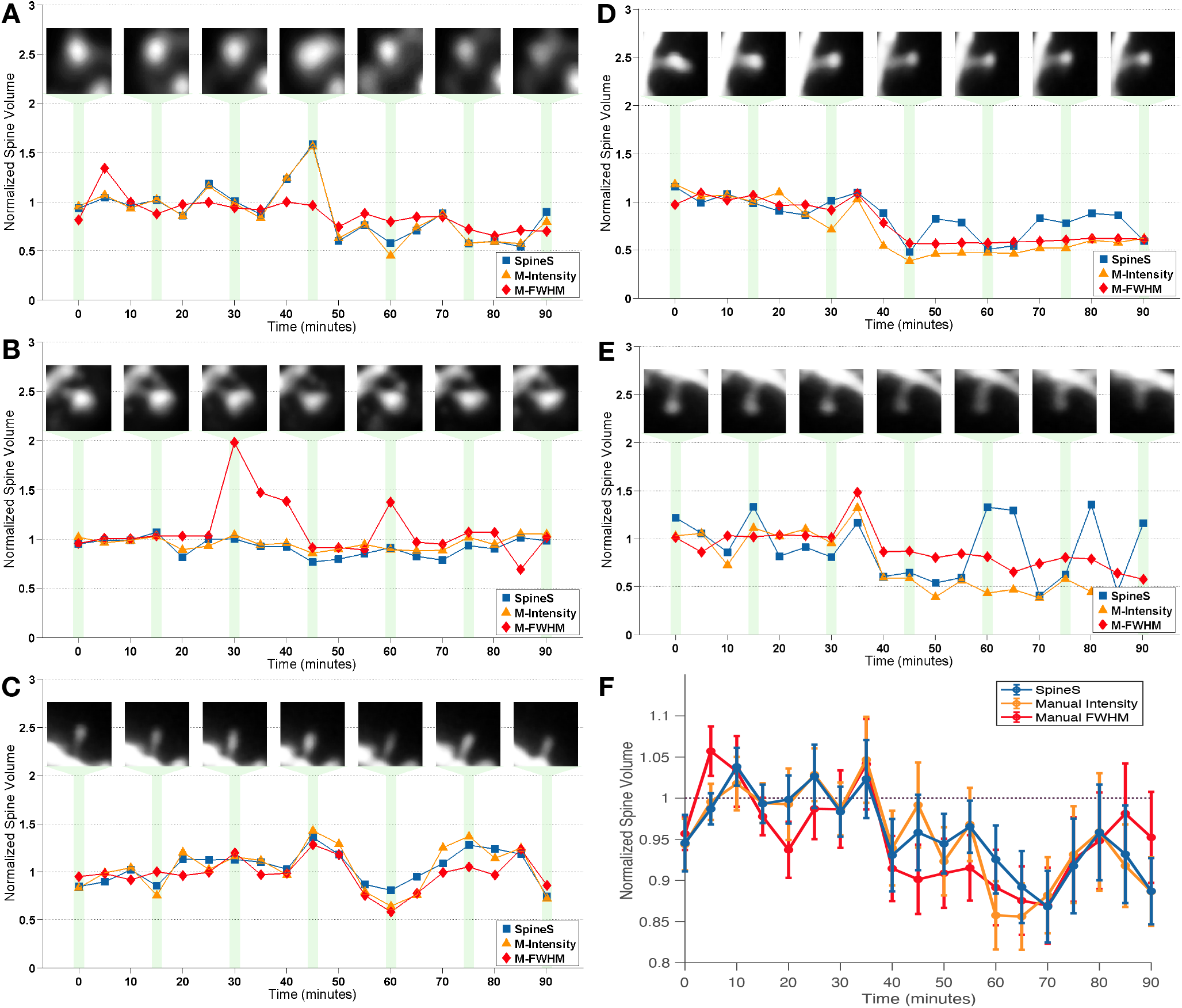
Comparison of automatic segmentation with manual segmentation and manual FWHM based volume estimation methods. Five examples from a chemical LTD experiment shown in panels A to E. Chemical agent DHPG is applied between 6th and 7th time points. (A) *SS_S–MI_* = 97.09, *SS_S–MF_* = 83.85, *SS_MI–MF_* = 84.23, (B) *SS_S–MI_* = 94.34, *SS_S–MF_* = 78.81, *SS_MI–MF_* = 81.16, (C) *SS_S–MI_* = 91.87, *SS_S–MF_* = 87.78, *SS_MI–MF_* = 88.16, (D) *SS_S–MI_* = 83.85, *SS_S–MF_* = 87.35, *SS_MI–MF_* = 89.86, (E) *SS_S–MI_* = 72.66, *SS_S–MF_* = 69.78, *SS_MI–MF_* = 77.61, (F) Average of 27 spines. SpineS: IFI based volume using automatic segmentations; Manual Intensity: IFI based volume using manual segmentations by an expert; Manual FWHM: FWHM based volume quantified by a different expert. Comparisons for all 27 spines can be found in Supplementary Table 1.

SpineS reports similar spine head volume results to the volume estimation obtained from manually segmented spine heads and manually quantified FWHM based volume estimation results by experts. As explained above, the FWHM based approach might lead to erroneous estimations if the spine head shape diverges from being spherical. The Integrated Fluorescence Intensity approach gives an arbitrary volume unit, which is sufficient when the relative volume change is the focus of the study. If real units of volumes are required, the user should perform a manual FWHM procedure for a number of spines to find a conversion factor into *μm*^3^ for the arbitrary IFI volume units.

### Performance on single spine stimulation plasticity data

One important application area of any spine analysis software is spine head volume and neck length tracking of single dendritic spines. In order to see the performance of our software in such experimental conditions, we collected images before and after the induction of LTD at a single spine using glutamate uncaging. Figure 7A shows the dendrite of interest before the stimulation. An LTD inducing glutamate uncaging stimulation was delivered before the 6*^th^* time point. Figure 7B shows the ROI of the stimulated spine over time. Figure 7C and D show the persistent shrinkage of the stimulated spine after stimulation. This form of stimulation does not appear to affect spine neck length (see. Figure 7D-below).

**Figure 7.**
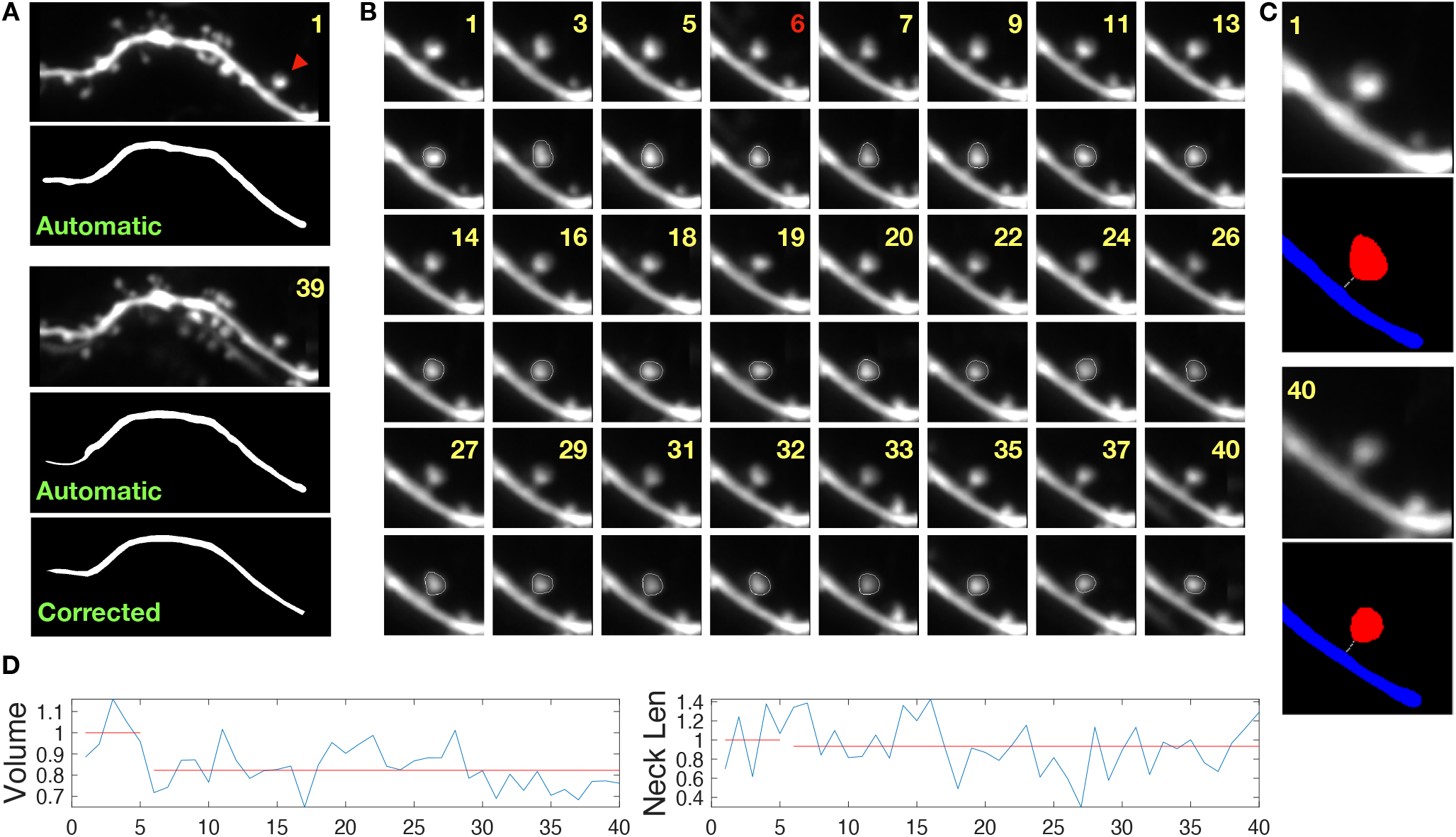
Glutamate uncaging induced single spine LTD experiment. (A) Dendritic segment before (upper two) and after stimulation (lower three). Stimulation was performed at the 6th time point. At the first time point the dendrite segmentation is perfect but at a later time-point, some fluorescence error leads to a problematic segmentation which is later corrected by manual interaction. The red arrowhead indicates the stimulated spine. (B) Spine head segmentation for 40 consecutive time points. We interleaved segmented and unsegmented rows to give a comparative sense of how the spines look for both representations. (C) Spine before (upper two) and after (lower two) undergoing LTD. Median filtered images are on the left, segmented images with neck paths are on the right (Blue: dendrite, Red: spine head, White: spine neck) (D) Normalized volume and spine neck results. (Blue: normalized trends, Red: linear fits for before and after stimulation.)

## Datasets

We tested our software on multiple datasets acquired from 4 different laboratories. Figures 4, 5 and 6 were produced using Dataset 1, Figure 7 using Dataset 2. Supplementary Figures 4 and 5 were the analysis results of Datasets 3 and 4, respectively. Acquisition summary of these datasets is shown in Table 1 and details are below.

**Table 1.**
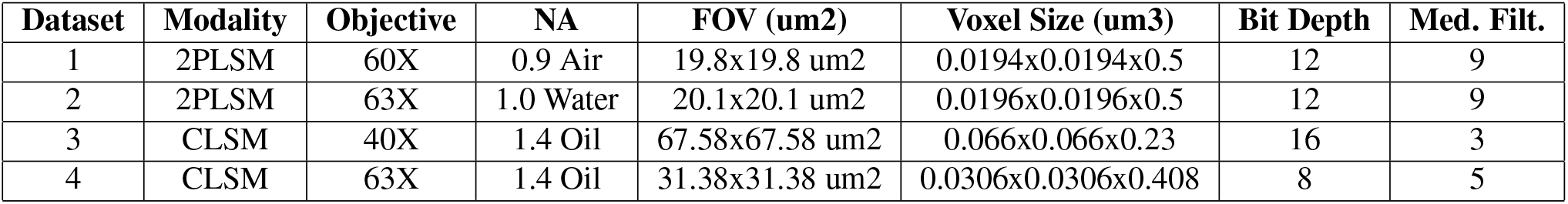
Details of the datasets used in the study. The last column presents the median filter numbers we used during the analysis that can be used as a guideline by the users.

| Dataset | Modality | Objective | NA | FOV (um2) | Voxel Size (um3) | Bit Depth | Med. Filt. |
| --- | --- | --- | --- | --- | --- | --- | --- |
| 1 | 2PLSM | 60X | 0.9 Air | 19.8x19.8 um2 | 0.0194x0.0194x0.5 | 12 | 9 |
| 2 | 2PLSM | 63X | 1.0 Water | 20.1x20.1 um2 | 0.0196x0.0196x0.5 | 12 | 9 |
| 3 | CLSM | 40X | 1.4 Oil | 67.58x67.58 um2 | 0.066x0.066x0.23 | 16 | 3 |
| 4 | CLSM | 63X | 1.4 Oil | 31.38x31.38 um2 | 0.0306x0.0306x0.408 | 8 | 5 |

### Dataset 1

Hippocampal neurons from mouse organotypic slice cultures postnatal day 7-9 were transfected using biolistic gene transfer with gold beads (10 mg, 1.6 um diameter, Biorad) coated with Dendra-2 (Evrogen) plasmid DNA (100*μ*g) or AFP using a Biorad Helios gene gun after 6 or 7 days in-vitro (DIV). Imaging experiments were performed 2 to 5 days post-transfection. Slices were perfused with artificial cerebrospinal fluid (ACSF) containing 127 mM *NaCl*, 2.5 mM *KCl*, 25 mM *NaHCO_3_*, 1.25 mM *NaH_2_PO*_4_, 25 mM D-glucose, 2 mM *CaCl_2_* and 1 mM *MgCl_2_* (equilibrated with O2 95%, CO2 5%) at room temperature at a rate of 1.5 ml/min. Two-photon imaging was performed using a galvanometer-based scanning system (Prairie Technologies, acquired by Bruker) on an Olympus BX61WI equipped with a 60X water immersion objective (0.9 NA), using a Ti:sapphire laser (910nm for imaging Dendra; Coherent) controlled by PrairieView software. Z-stacks (0.5*μ*m axial spacing) from secondary or tertiary dendrites from CA1 neurons were collected every 5 min for up to 4 hours. 1024×1024 pixel, xy pixel size = 0.0193*μ*m, FOV 19.8×19.8*μ*m.

### Dataset 2

Cultured hippocampal slices were prepared from postnatal day 7-10 mice SHANK3 (Jax, no. 017889), as described^42^. The cultures were transfected by Gene gun (Bio Rad) with AFP after 3-6 days in vitro (DIV). Slices were perfused with ACSF containing 127 mM *NaCl*, 2.5 mM KCl, 25 mM *NaHCO_3_*, 1.25 mM *NaH_2_PO_4_*, 25 mM D-glucose, 2 mM *CaCl_2_* and 1 mM *MgCl_2_* (equilibrated with O2 95% CO2 5%) at room temperature at a rate of 1.5 ml/min. Two-photon imaging was performed using a LSM 710 microscope (Zeiss) based on galvanometer scanning system controlled by Zen black software, equipped with W Plan Apochromat 63X water immersion objective (1.0 NA). The light source was a Ti:Sapphire laser (Chameleon Ultra II, Coherent) tuned to 910 nm for imaging and 720nm for uncaging experiments controlled by Zen black software. Z-stacks (0.3*μ*m axial spacing) from secondary or tertiary dendrites from CA1 neurons were collected every 5 min up to 4 hours. For uncaging LTD experiments the laser was positioned at 0.5*μ*m from the center of spine, then light pulses were delivered at low frequency to induce LTD.

### Dataset 3

Hippocampal organotypic cultures were made from P6/7 wistar rats and transfected using biolistic gene transfer with gold beads coated with mCherry. Images were acquired every 10min at DIV9 from a secondary dendrite of a CA1 pyramidal neuron. Confocal image stacks were collected using a 40x objective with 4X zoom.

### Dataset 4

Hippocampal slices from wistar rats at postnatal day 7 were transfected with CMV-eGFP 4 days before imaging. Imaging was done at day in vitro 15. Images were acquired at intervals of 5 mins. The microscope was a Zeiss LSM 710 AxioObserver, 63X oil immersion objective (1.4 NA), Emission wavelength 574, excitation 488, EGFP, pixel size 0.0306 × 0.0306 × 0.0408*μ*m, zoom 4.3.

## Discussion

We present an image-processing tool for longitudinal quantification of dendritic spine features. The proposed tool yields good results in terms of accuracy and run times for spine analysis. Results demonstrate that IFI and FWHM can be used interchangeably for individual volume trend assessment for cases in which the spine maintains a regular spherical shape throughout all the time points, or could be used interchangeably when pooled data is the key to research. SpineS provides a means for post quality assessment which gives users the flexibility to reject or correct individual spine segmentations and neck paths for any time point. This feature provides an important quality control point of the software, as segmentation quality is subject to fluorescence intensity differences between the spine head and neck, which can vary. Overall, the SpineS toolbox improves the speed of image analysis, reducing analysis time from days to hours, while ensuring the quality of the analysis. Another advantage of SpineS is that it provides a less subjective quantification of structure. Manual FWHM measurements require the user to orient and place a line through the spine head to determine the center of the spine in each image which can introduce experimental bias.

As a future step, we are currently developing a deep learning-based segmentation tool to embed into the current SpineS software to make the dendrite and spine segmentation steps faster and more accurate. We aim to translate the entire code into Python with QT framework to increase the availability.

### Data and Code Availability

Data and code can be downloaded from https://github.com/argunsah/SpineS.

## Acknowledgements

We thank Ana Luisa Carvalho for datasets 3 and 4, Débora Serrenho for testing the software and Asim Iqbal for reading the manuscript and providing feedback. A.Ö.A. was partially supported by Fundação para a Ciência e a Technologia (FCT) grant SFRH /BD/51264/2010. E.E. was supported by Personalized Health and Related Technologies (PHRT), project number 222, ETH domain. A.F.H. was supported by FCT grant SFRH/BD/51265/2010. The authors would also like to acknowledge the financial support of the European Research Council (ERC) via a Starting Grant to T.K (no. 679175), Fundação Bial (161/10-2010) and (PTDC/SAU-NMC/122035/2010) to I.I., Consejo Nacional de Ciencia y Tecnología to Y.R.C. (254878) and Scientific and Technological Research Council of Turkey to D.Ü. and M.Ç. (113E603).

## Author contributions

A.Ö.A, I.I. and D.Ü. conceived the study. A.Ö.A, E.E., M.U.G. and D.Ü. wrote the code. A.Ö.A, Y.R.C. and A.F.H. conducted the experiments. A.Ö.A. made the GUI and analysed the results. A.Ö.A, E.E. and D.Ü. wrote the manuscript. T.K., M.C. I.I. and D.Ü. provided funding.

