## Supplementary Material for "SpineS: An interactive time-series analysis software for dendritic spines"

[1]Champalimaud Research, Champalimaud Centre for the Unknown, 1400-038 Lisbon, Portugal
[2]Laboratory of Neural Circuit Assembly, Brain Research Institute (HiFo), University of Zürich, Zürich, Switzerland.
[3]Neuroscience Center Zurich (ZNZ), UZH/ETH Zürich, Zürich, Switzerland.
[4]Computer Vision Laboratory, ETH Zürich, Zürich, Switzerland.
[5]Signal Processing and Information Systems Laboratory, Sabancı University, Tuzla, İstanbul 34956, Turkey.
[6]Department of Electrical and Computer Engineering, Boston University, Boston, MA 02215, USA.
[7]Departamento de Neurodesarrollo y Fisiología, Instituto de Fisiología Celular, Universidad Nacional Autónoma de México, Ciudad de México, C.P. 04510, Mexico.
[8]Department of Electrical and Computer Engineering, University of Rochester, Rochester, NY 14627, USA.
[9]Department of Pathology and Cell Biology, Columbia University, New York, NY 10032, USA.
[10]Department of Biomedical Engineering, İzmir University of Economics, İzmir, Turkey.
[11]Department of Electrical and Electronics Engineering, İzmir Demokrasi University, İzmir, Turkey.
[+]Equal Contribution

### ABSTRACT

Live fluorescence imaging has shown the dynamic nature of dendritic spines, with changes in shape occurring both during development and in response to activity. The structure of a dendritic spine positively correlates with its functional efficacy. Learning and memory studies have shown that great deal of the information stored by a neuron is contained in the synapses. High precision tracking of synaptic structures can give hints about the dynamic nature of memory and help us to understand how memories evolve both in biological and artificial neural networks. Experiments that aim to investigate the dynamics behind the structural changes of dendritic spines require the collection and analysis of large time-series datasets. In this paper, we present an open-source software called SpineS for the automatic longitudinal structural analysis of dendritic spines with additional features for manual intervention to ensure optimal analysis. Our extensive experimental analyses on multiple datasets demonstrate that SpineS can achieve a high-level performance on samples collected both by two-photon and confocal imaging systems.

### Supplementary Information

| Dendrite | Spine | SpineS vs Manual Intensity | SpineS vs Manual FWHM | Manual Intensity vs Manual FWHM |
|---|---|---|---|---|
| 1 | 1 | 97.52 | 86.34 | 87.01 |
| | 2 | 88.73 | 80.23 | 84.71 |
| | 3 | 91.23 | 91.12 | 88.68 |
| | 4 | 94.59 | 74.72 | 72.93 |
| 2 | 5 | 97.09 | 83.85 | 84.23 |
| | 6 | 88.70 | 87.4 | 90.15 |
| | 7 | 90.81 | 89.06 | 88.01 |
| 3 | 8 | 95.88 | 90.01 | 88.31 |
| | 9 | 90.80 | 82.33 | 85.52 |
| | 10 | 97.08 | 79.10 | 80.44 |
| | 11 | 86.01 | 90.06 | 86.27 |
| | 12 | 87.75 | 78.34 | 74.54 |
| | 13 | 94.34 | 78.81 | 81.16 |
| 4 | 14 | 88.22 | 89.12 | 87.24 |
| | 15 | 83.85 | 87.35 | 89.86 |
| 5 | 16 | 89.37 | 82.26 | 82.27 |
| | 17 | 90.33 | 89.39 | 87.98 |
| 6 | 18 | 91.87 | 87.78 | 88.19 |
| | 19 | 72.66 | 69.78 | 77.61 |
| | 20 | 83.63 | 84.67 | 88.86 |
| 7 | 21 | 94.03 | 92.66 | 95.14 |
| | 22 | 78.77 | 71.41 | 91.36 |
| 8 | 23 | 94.09 | 92.32 | 93.28 |
| | 24 | 94.56 | 91.14 | 90.74 |
| 9 | 25 | 86.42 | 84.32 | 86.00 |
| | 26 | 96.07 | 67.95 | 70.77 |
| | 27 | 93.08 | 84.62 | 81.52 |
| Mean | | 90.28 | 83.93 | 85.29 |
| S.D. | | 5.83 | 6.93 | 6.01 |

**Table 1.** Comparison of automatic segmentation with manual segmentation and manual FWHM based volume estimation methods. Comparisons for all 27 spines. SpineS: IFI based volume using automatic segmentations; Manual Intensity: IFI based volume using manual segmentations by an expert; Manual FWHM: FWHM based volume quantified by a different expert.(Dataset 1)

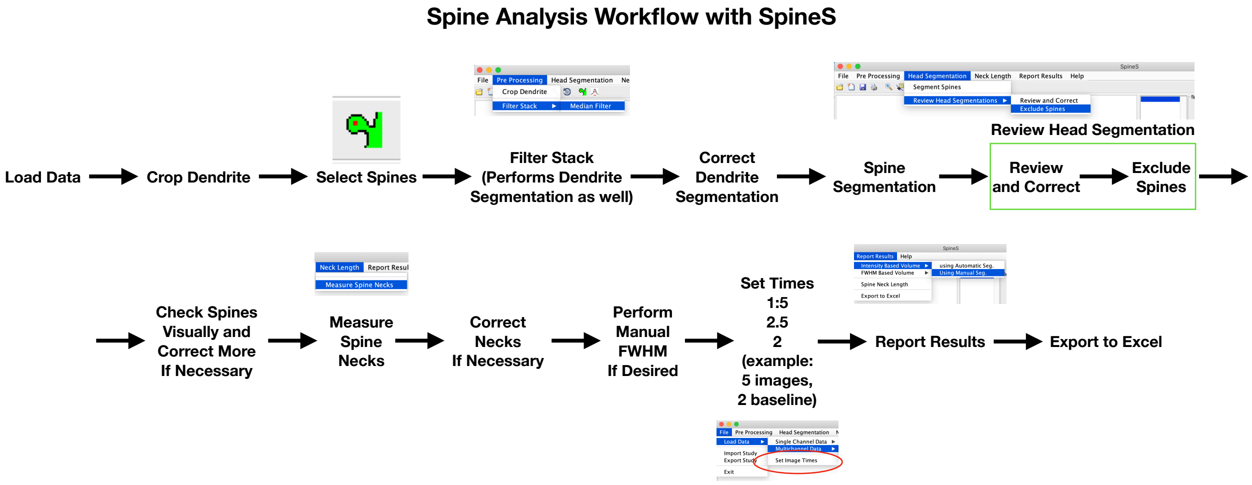


**Figure 1.** Steps to follow to analyse a dataset.

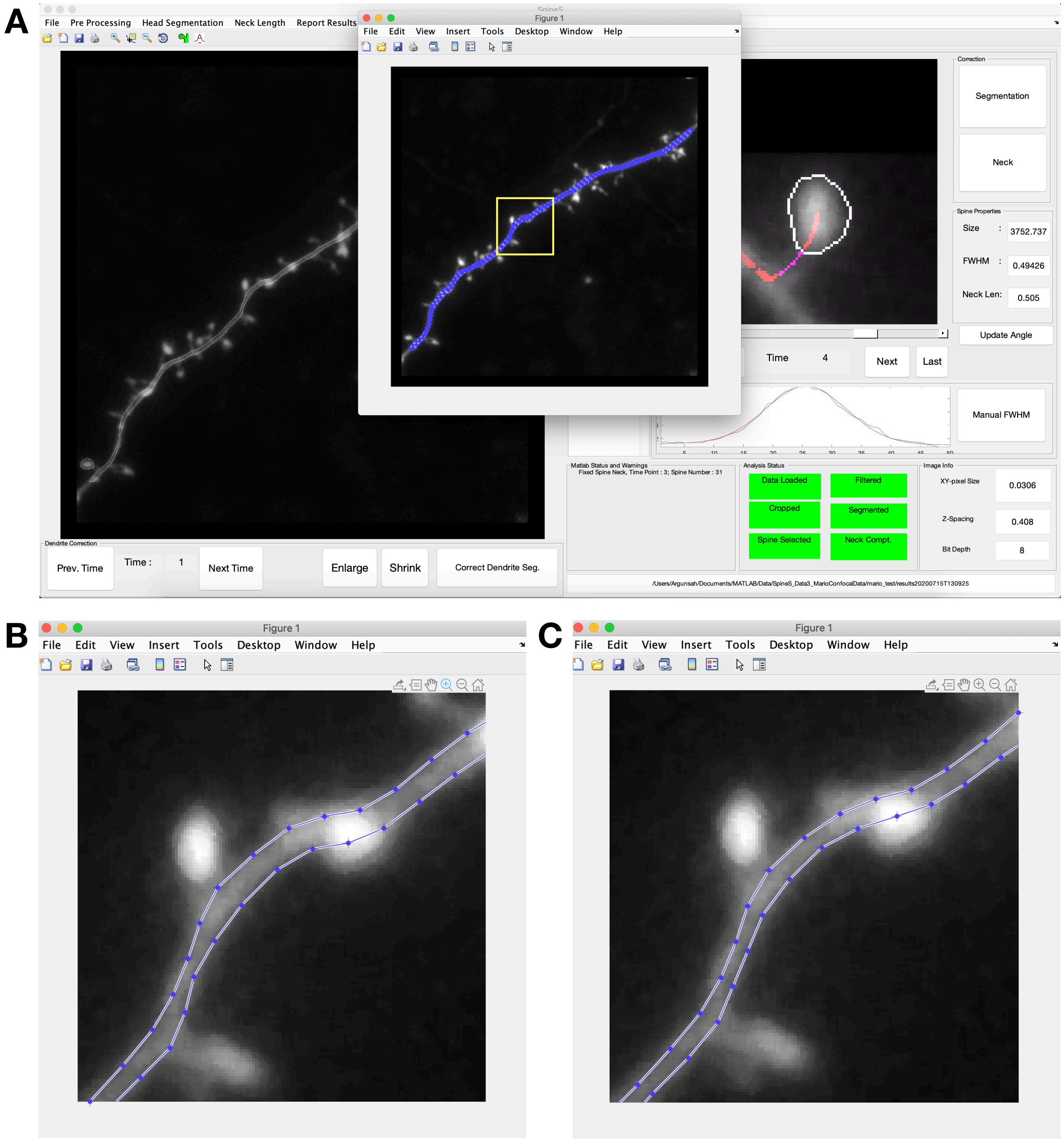


**Figure 2.** Dendrite segmentation correction. (A) *Correct Dendrite Seg.* button opens a new window with segmented dendrite in discreet movable points. User should move imperfect points by holding the left mouse button and dragging. Double click at one of the points saves the segmentation. (B) Example of an imperfect segmentation. (C) Segmented dendrite after manual intervention.

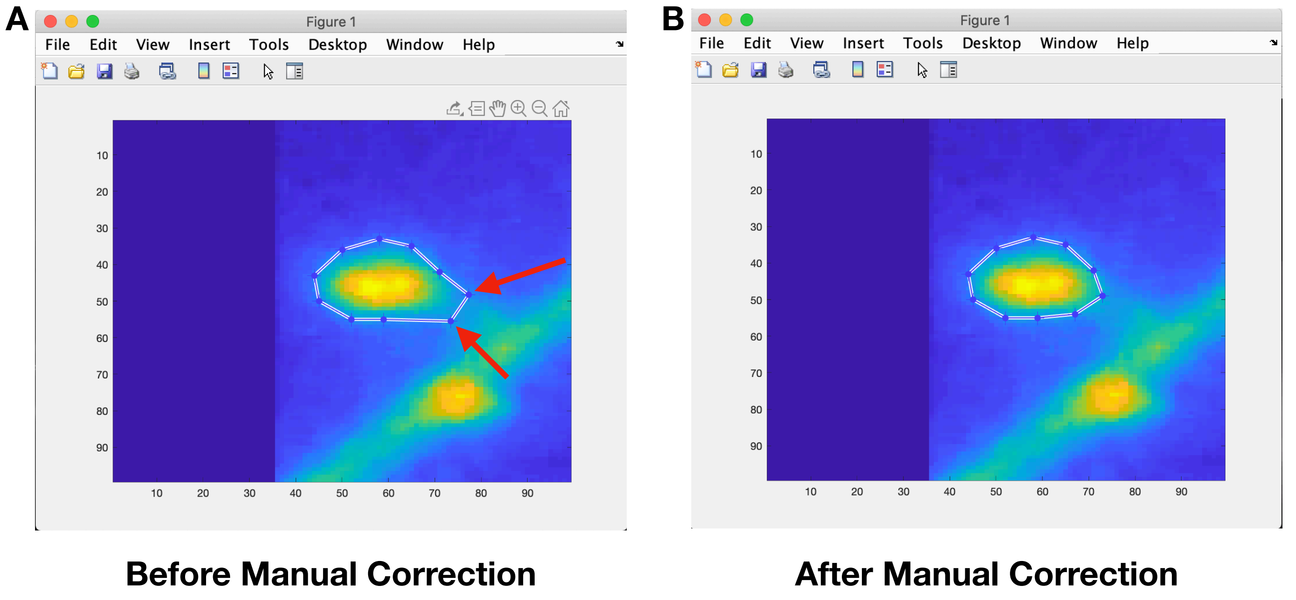


**Figure 3.** Spine segmentation correction. *Segmentation* button under Correction panel opens a new window with segmented spine in discreet movable points. User should move imperfect points by holding the left mouse button and dragging. Double click at one of the points saves the segmentation. (A) Example of an imperfect segmentation. (B) Segmented spine head after manual correction.

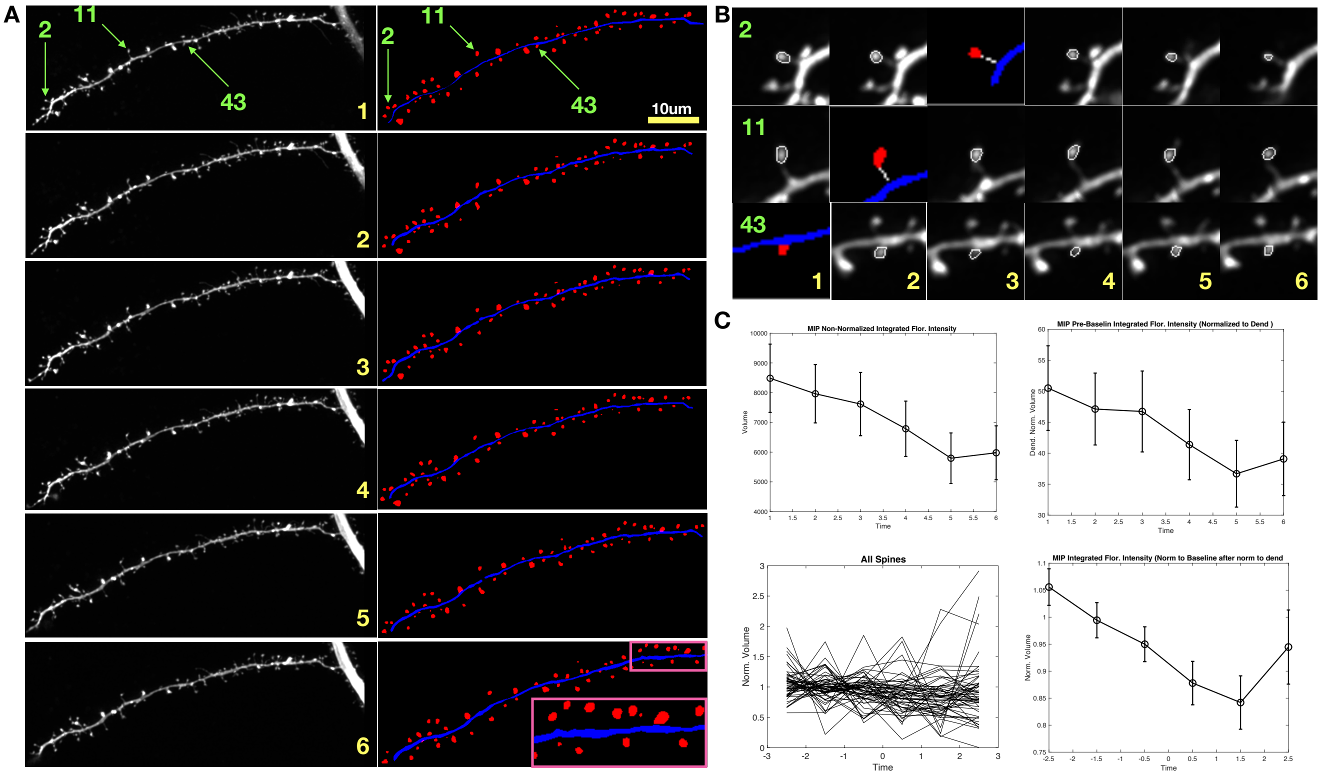


**Figure 4.** Dataset 3 Analysis Results. 57 spines were analyzed at 6 consecutive time points. (A) Images of the dendrite on the left column, segmented dendrite and spines on the right. (B) Three example spines and their segmentations and/or neck paths. (C) Results of the spine head volumes before and after normalization on average and individually. Yellow numbers represent time, green numbers represent spines.

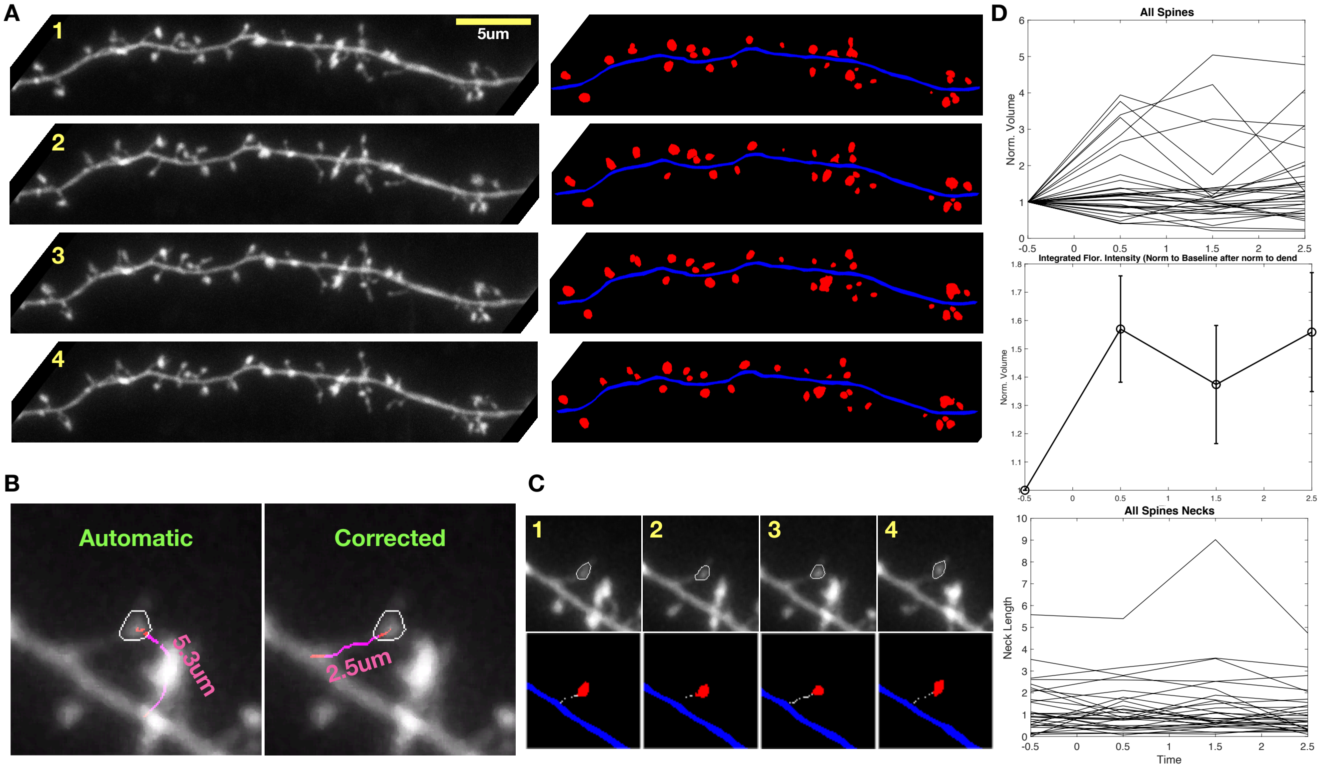


**Figure 5.** Dataset 4 Analysis Results. 36 spines were analyzed at 4 consecutive time points. (A) Images of the dendrite on the left column, segmented dendrite and spines on the right. (B) An example of a bad spine neck path on the left and after manual correction on the right. Spine neck is traced from the center of the spine to the center of the dendrite (red path) but neck length is computer from the edges of spine and dendrite segmentations (magenta path). (C) Example of a segmented spine on top, segmentation as well as neck path on the bottom. (D) Top: Individual spine normalized volume over time, Middle: Average normalized volume over time. Bottom: Individual spine neck lengths. Yellow numbers represent time, green number represent spines.
